# DNA Repair Gene Expression Adjusted by the PCNA Metagene Predicts Survival in Multiple Cancers

**DOI:** 10.1101/446377

**Authors:** Leif E. Peterson, Tatiana Kovyrshina

**Affiliations:** Department of Healthcare Policy and Research, Weill Cornell Medical College, Cornell University, New York City, New York 10065 USA; Center for Biostatistics, Institute for Academic Medicine, Houston Methodist Research Institute, 6565 Fannin Street, Houston, Texas 77030 USA

## Abstract

One of the hallmarks of cancer is the existence of a high mutational load in driver genes, which is balanced by upregulation (downregulation) of DNA repair pathways, since almost complete DNA repair is required for mitosis. The prediction of cancer survival with gene expression has been investigated by many groups, however, results of a comprehensive re-evaluation of the original data adjusted by the *PCNA* metagene indicate that only a small proportion of genes are truly predictive of survival. However, little is known regarding the effect of the *PCNA* metagene on survival prediction specifically by DNA repair genes. We investigated prediction of overall survival (OS) in 18 cancers by using normalized RNA-Seq data for 126 DNA repair genes with expression available in TCGA. Transformations for normality and adjustments for age at diagnosis, stage, and *PCNA* metagene were performed for all DNA repair genes. We also analyzed genomic event rates (GER) for somatic mutations, deletions, and amplification in driver genes and DNA repair genes. After performing empirical p-value testing with use of randomly selected gene sets, it was observed that OS could be predicted significantly by sets of DNA repair genes for 61% (11/18) of the cancers. Pathway activation analysis indicates that in the presence of dysfunctional driver genes, the initial damage signaling and minor single-gene repair mechanisms may be abrogated, but with later pathway genes fully activated and intact. Neither *PARP1* or *PARP2* were significant predictors of survival for any of the 11 cancers. Results from cluster analysis of GERs indicates that the most opportunistic set of cancers warranting further study are AML, colorectal, and renal papillary, because of their lower GERs for mutations, deletions, and amplifications in DNA repair genes. However, the most opportunistic cancer to study is likely to be AML, since it showed the lowest GERs for mutations, deletions, and amplifications, suggesting that DNA repair pathway activation in AML is intact and unaltered genomically. In conclusion, our hypothesis-driven focus to target DNA repair gene expression adjusted for the *PCNA* metagene as a means of predicting OS in various cancers resulted in statistically significant sets of genes.

**Author summary:** The proliferating cell nuclear antigen (*PCNA*) protein is a homotrimer and activator of polymerase *δ*, which encircles DNA during transcription to recruit other proteins involved in replication and repair. In tumor cells, expression of *PCNA* is highly upregulated; however, *PCNA*-related activity is a normal process for DNA transcription in eukaryotes and therefore is not considered to play a central role in the selective genetic pressure associated with tumor development. Since *PCNA* is widely co-regulated with other genes in normal tissues, we developed workflow involving several functional transforms and regression models to “remove” the co-regulatory effect of *PCNA* on expression of DNA repair genes, and predicted overall cancer survival using DNA repair gene expression with and without removal of the *PCNA* effect. Other adjustments to survival prediction were employed, such as subject age at diagnosis and tumor stage. Random selection of gene sets was also employed for empirical p-value testing to determine the strength of the *PCNA* effect on DNA repair and overall survival adjustments. Since TCGA RNA-Seq data were used, we also characterized the frequency of deletions, amplifications, and somatic mutations in the DNA repair genes considered in order to observe which genomic events are the most frequent for the cancers evaluated.

## 1 INTRODUCTION

Recent developments in cancer research based on next-generation sequencing technology indicates that a hallmark of sporadic cancers is that they exhibit a lifelong accumulation of pathogenic somatic mutations in “driver genes,” otherwise known as tumor suppressor genes and oncogenes [1]. There are many different biological activities and molecular functions of driver genes, including, for example, DNA binding, protein binding, transcription factor activity, receptor activity, magnesium and calcium ion binding, actin binding, protein kinase activity, etc. Altogether, most cancer driver genes showing accumulation of somatic mutations are commonly not involved in DNA repair and mitosis, which involve genes that are functionally related to repair of DNA single-strand breaks, double-strand breaks, and activation of cell-cycle checkpoints. It is clear that upregulated DNA repair within tumor cells helps reduce the mutational load caused by the buildup of somatic mutations conferring a selection advantage for mitosis, leading to clonal expansion and cellular growth.

The *PCNA* protein is a ring-like molecule that serves as a co-factor for polymerase *δ*, and surrounds DNA during strand synthesis to recruit proteins needed for DNA replication [2]. In a previous investigation by Ge et al. [3], 131 mRNAs were identified in 36 types of normal tissues whose expression correlated *r* > 0.65 with expression of *PCNA*. *PCNA* by itself is not a tumor suppressor gene or oncogene and is merely a gene whose expression is upregulated during DNA replication. Venet et al. [4] replicated 47 breast cancer gene profile (survival) studies using the same gene expression data used in the original studies, and found that 28 profiles (60%) were less significant than randomly selecting (1000 times) the same number of genes reported in each profile. In addition, after collapsing expression of the 131 *PCNA*-associated genes into a median (called *PCNA* “metagene”), it was observed that 91% of the genes predictive of survival were significantly correlated with the *PCNA* metagene, since nearly 50% of genome is associated with *PCNA* [4]. Results of their investigation revealed that the majority of published gene sets which were reported to predict survival significantly (in breast cancer) perform worse than randomly selecting the same number of genes from the entire expression sets, and that removing the effect of the *PCNA* metagene from expression of the original gene sets resulted in much worse survival prediction.

In this investigation, we specifically focused on DNA repair genes, under the hypothesis that the adjustment of DNA repair gene expression by use of the *PCNA* metagene would result in a significant association with survival in 18 cancers for which data exist in the TCGA [5]. We also hypothesized that patterns of somatic mutations, deletions, and amplifications in cancer-specific driver genes and the DNA repair genes considered would provide new insight into the patterns of genomic alteration observed in tumor cells. Results of the computational analyses were used for generating lists of DNA repair genes, whose downregulation or upregulation therapeutically may potentially confer prolonged survival of patients. As such, the focus of this project was to identify patterns of DNA repair gene expression for recommending expression exploitation for potential transgenics, knock-down, or overexpression as a means of cancer therapy.

## 2 DATA

### Cancer data

The data used in this investigation were derived from nextgen sequencing of tumors in the The Cancer Genome Atlas (TCGA) [5]. We investigated DNA repair gene expression in 18 cancers for which nextgen sequencing and RNA-seq expression data were available from cBio-Portal (http://www.cbioportal.org) [6,7]. Specifically, the cancers for which only age at diagnosis was available, and not pathological stage, in the TCGA data were acute myelogenous leukemia (AML), bladder, low grade gliomas, gliobastoma multiforme (GBM), head and neck, and sarcoma. Cancers for which both age at diagnosis and pathological stage were available included breast, cervical, colorectal, liver, lung, lung squamous cell, melanoma, ovarian, renal clear cell, renal papillary, stomach, and uterine. Altogether, this resulted in a total of 18 cancers that were considered.

### Cancer-specific consensus driver genes

We obtained a consensus list of the top 20 driver genes for each cancer considered from the DriverDB database [8], based on identification by at least 2 tools (default) for each cancer, since requesting a higher consensus could result in fewer than 20 driver genes for some cancers. DriverDB assembles together lists of the top ranked driver genes determined from the use of 15 packages, including ActiveDriver, Dendrix, MDPFinder, Simon, NetBox, OncodriveFM, MutSigCV, MEMo, CoMDP, DawnRank, DriverNet, e-Driver, iPAC, MSEA, and OncodriveCLUST. A list of the top 20 driver genes used for each cancer has been previously reported [9].

### Expression, mutations, deletions, and amplifications

RNA-Seq based normalized expression values for and somatic mutations in DNA repair genes were also obtained from cBio-Portal [6,7]. We also acquired high-confidence deletions and amplifications from cBio-Portal, where a deletion was defined as full homozygous loss with a GISTIC score [10] of −2, and an amplification was defined as high-level gain with a GISTIC score of 2. Low-level deletions (heterozygous loss) and low-level gain (low-level amplifications) with GISTIC scores of −1 and 1, respectively, were not used.

### Direct Reversal Repair, DRR

Direct reversal DNA repair (DRR) is a single step reaction of removal of the methyl- or photoadducts. DRR is provided to methylphosphotriesters (direct removal of the alkyl damage by nucleophilic Cys residues), alkyltransferases (repair of O6-alkylguanine by *MGMT*), oxidative dealkylation (*ALKB* proteins), and photolyase (direct reversal of the thymine dimer created by UV light in CTD photolyase and 6-4TT photolyase). The DRR genes used in this study were: *ALKBH2*, *ALKBH3*, and *MGMT*.

### Base Excision Repair, BER

Base excision DNA repair (BER) corrects base lesions generated by oxidative, alkylation, deamination, and depurinatiation/depyrimidination damage. BER is initiated by DNA glycosylases, which recognize and catalyze removal of damaged bases. Downstream enzymes carry out strand incision, gap-filling, and ligation. BER involves two general pathways: short-patch (SP-BER) and long-patch (LP-BER). BER genes employed in this study included: *APEX1*, *APEX2*, *APTX*, *FEN1*, *HUS1*, *LIG1*, *LIG3*, *MBD4*, *MPG*, *MUTYH*, *NEIL1*, *NEIL2*, *NEIL3*, *NTH*, *OGG1*, *PARP1*, *PARP2*, *PCNA*, *PNKP*, *POLB*, *POLD1*, *POLE*, *POLH*, *POLL*, *RECQL2*, *SMUG1*, *TDG*, *UNG*, and *XRCC1*.

### Non-homologous End-joining, NHEJ

Non-homologous end-joining (NHEJ) repairs DSBs at all stages of the cell-cycle, bringing about the ligation of two double-strand breaks (DSBs) without the need for sequence homology, and therefore NHEJ is error-prone. NHEJ is referred to as “non-homologous” because the break ends are directly ligated without the need for a homologous template, in contrast to homologous recombination, which requires a homologous sequence to guide repair. NHEJ typically utilizes short homologous DNA sequences called microhomologies, which are often present in single-stranded overhangs on the ends of DSBs. Inappropriate NHEJ can lead to translocations and telomere fusion, which are common hallmarks of tumor cells. The NHEJ genes considered in this investigation were: *DCLRE1C*, *XRCC6*, *XRCC5*, *LIG4*, *NHEJ1*, *POLL*, *POLM*, *PRKDC*, *RECQL2*, *XPF*, and *XRCC4*.

### Mismatch Repair, MMR

DNA mismatch repair (MMR) is responsible for correction of replication errors (mismatches and small insertions and deletions) that escape the proofreading activity of a DNA polymerase. MMR is initiated by two proteins homologous to MutS and MutL: MutS*α* and MutL*α*. Mutations in the genes coding MutS and MutL homologs have been linked with the Lynch syndrome, which is characterized by an increased risk of developing cancer. MMR genes included in this study were: *EXO1*, *LIG1*, *MLH1*, *MLH3*, *MSH2*, *MSH3*, *MSH6*, *PMS1*, *PMS2*, and *POLD1*.

### Translesion Synthesis, TLS

Translesion synthesis (TLS) is a process involving specialized DNA polymerases which replicate across from DNA lesions. TLS aids in resistance to DNA damage, presumably by restarting stalled replication forks or filling in gaps that remain in the genome due to the presence of DNA lesions. TLS has the potential to produce mutations. The TLS genes considered in this study were: *POLH*, *POLI*, *POLK*, *POLM*, *POLN*, *POLQ*, *REV1*, and *REV3L*.

### DNA Damage Signaling, DDS

DNA damage induces several cellular responses including DNA repair, cell-cycle checkpoint activity, and triggering of apoptotic pathways. DNA damage checkpoints are associated with biochemical pathways that end delay or arrest of cell-cycle progression. Such checkpoints engage damage sensor proteins, such as the *RAD9-RAD1-HUS1* (9-1-1) complex, and the *RAD17-RFC* complex, in the detection of DNA damage and transduction of signals to *ATM*, *ATR*, *CHK1* and *CHK2* kinases. In addition, *CHK1* and *CHK2* kinases regulate *CDC25*, p21 and p53 that ultimately inactivate cyclin-dependent kinases (CDKs), which inhibit cell-cycle progression. The DDS genes considered included: *ATM*, *ATR*, *ATRIP*, *BLM*, *BRCA1*, *CCNH*, *CDK7*, *CDKN1A*, *CHEK1*, *CHEK2*, *COPS5*, *DCLRE1A*, *DCLRE1B*, *FANCA*, *FANCC*, *GPS1*, *HUS1*, *MDC1*, *MNAT1*, *MRE11A*, *NBN*, *RAD1*, *RAD17*, *RAD18*, *RAD23A*, *RAD50*, *RAD9A*, *RFC1*, *RFC2*, *RFC3*, *RFC4*, *RFC5*, *TOPBP1*, and *TP53*.

### Homologous Recombination Repair, HRR

Homologous recombination repair (HRR) is a type of genetic recombination in which nucleotide sequences are exchanged between two similar or identical molecules of DNA. HRR is an “error free” mechanism which acts on DSBs occurring within replicated DNA (replication-independent DSBs) or on DSBs that are generated at broken replication forks (replication-dependent DSBs). HRR also involves processing of the ends of the DNA double-strand break, homologous DNA pairing and strand exchange, repair DNA synthesis, and resolution of the heteroduplex molecules. The HRR genes used in this project included: *BLM*, *BRCA1*, *BRCA2*, *C19GRF40*, *EME1*, *EME2*, *FANCA*, *FANCB*, *FANCC*, *FANCD2*, *FANCE*, *FANCF*, *FANCG*, *FANCI*, *FANCL*, *MRE11A*, *MSH4*, *MSH5*, *MUS81*, *RAD51*, and *RAD52*.

### Nucleotide Excision Repair, NER

Nucleotide excision repair (NER) removes UV-induced damage (thymine dimers and 6-4-photoproducts) as well as other kinds of DNA damage, which produce bulky distortions in the shape of DNA double helix. NER enzymes recognize bulky distortions in the shape of the DNA double helix, and only repair damaged bases that can be removed by a specific glycosylase. Specifically, NER entails removal of a short single-stranded DNA segment that includes the lesion, creating a single-strand gap (20-30 nucleotides) in the DNA, which is subsequently filled in by DNA polymerase d or e by copying the undamaged strand. Polymorphisms in NER proteins include Xeroderma pigmentosum and Cockayne’s syndrome. The NER genes considered in this study included: *CSA*, *CSB*, *CUL4A*, *DDB1*, *DDB2*, *ERCC1*, *GTF2H1*, *GTF2H2*, *GTF2H3*, *GTF2H4*, *GTF2H5*, *LIG1*, *MMS19*, *PCNA*, *PGLD1*, *POLE*, *RAD23B*, *RFC1*, *RPA1*, *XPA*, *XPB*, *XPC*, *XPD*, *XPF*, and *XPG*.

## 3 METHODS

### PCNA Metagene

For each tumor (row), we obtained RNA-Seq derived normalized expression values of the 131 PCNA-related genes [4], and collapsed their expression values down to median expression, which is termed the “PCNA metagene.” Next, the *PCNA* metagene (median) and expression values for all of the DNA repair genes listed above were transformed into van der Waerden (VDW) scores. This transformation simply involved first transforming expression values for each gene into percentiles, and then substituting the percentiles as probabilities in the inverse standard normal function, i.e., *Z* = Φ^−1^(*ptile*), to obtain standard normal variates of expression, which are distributed with mean zero and variance unity.

### Maximum Likelihood (ML) Survival Prediction

VDW scores for each DNA repair genes were regressed separately on the VDW scores for age, VDW scores for tumor stage, and VDW scores for *PCNA* metagene, and the residuals were taken as the new DNA repair gene expression values. Residuals for each DNA repair gene were then binarized into (0,1) by splitting on the median, to form a grouping variable which was employed in Kaplan-Meier (KM) survival analysis with overall survival (OS) status. Each DNA repair gene which resulted in significant maximum likelihood (ML) estimates of the KM logrank tests, i.e., x^2^_(1)_) p-value < 0.05, was appended to a list of *p* genes. Eigendecomposition of the correlation matrix of the p significant genes was then performed with Varimax orthogonal rotation, and the principal components (PCs) for all *p* dimensions were extracted. Each PC was then transformed into a binary grouping variable for KM input, by assigning negative PC values to 0 and positive to 1. The single group-transformed PC which resulted in the greatest x^2^_(1)_ value during KM analysis was identified, and called the “best binarized PC.” For cancers without stage available in TCGA, the best binarized PC when age and *PCNA* were used for residual generation was input into Cox proportional hazards (PH) regression as a continuous variable (i.e., PC score) to determine whether positive values prolonged or shortened OS. Whereas, for cancers with stage available in TCGA, the best binarized PC when when age, stage, and *PCNA* were used for residual generation was input into Cox proportional hazards (PH) regression. For the best binarized PC under evaluation, if the Cox PH hazard ratio (*HR*) < 1, then genes having positive loading on the best binarized PC were beneficial to OS if upregulated, whereas genes whose loadings were negative were considered hazardous, and would need to be downregulated in order to be beneficial. Analogously, if the Cox PH hazard ratio (*HR*) > 1, it meant that positive PC values were deleterious, and therefore genes that loaded negatively on this PC would need to be upregulated to be beneficial, and genes that loaded positively on this PC would need to be downregulated in order to be beneficial to OS. Figure 1 illustrates the workflow employed, outlining the various steps used for establishing the best binarized PC for each cancer, and whether positive loadings on the best binarized PC prolonged or shortened OS.

**Fig 1.**
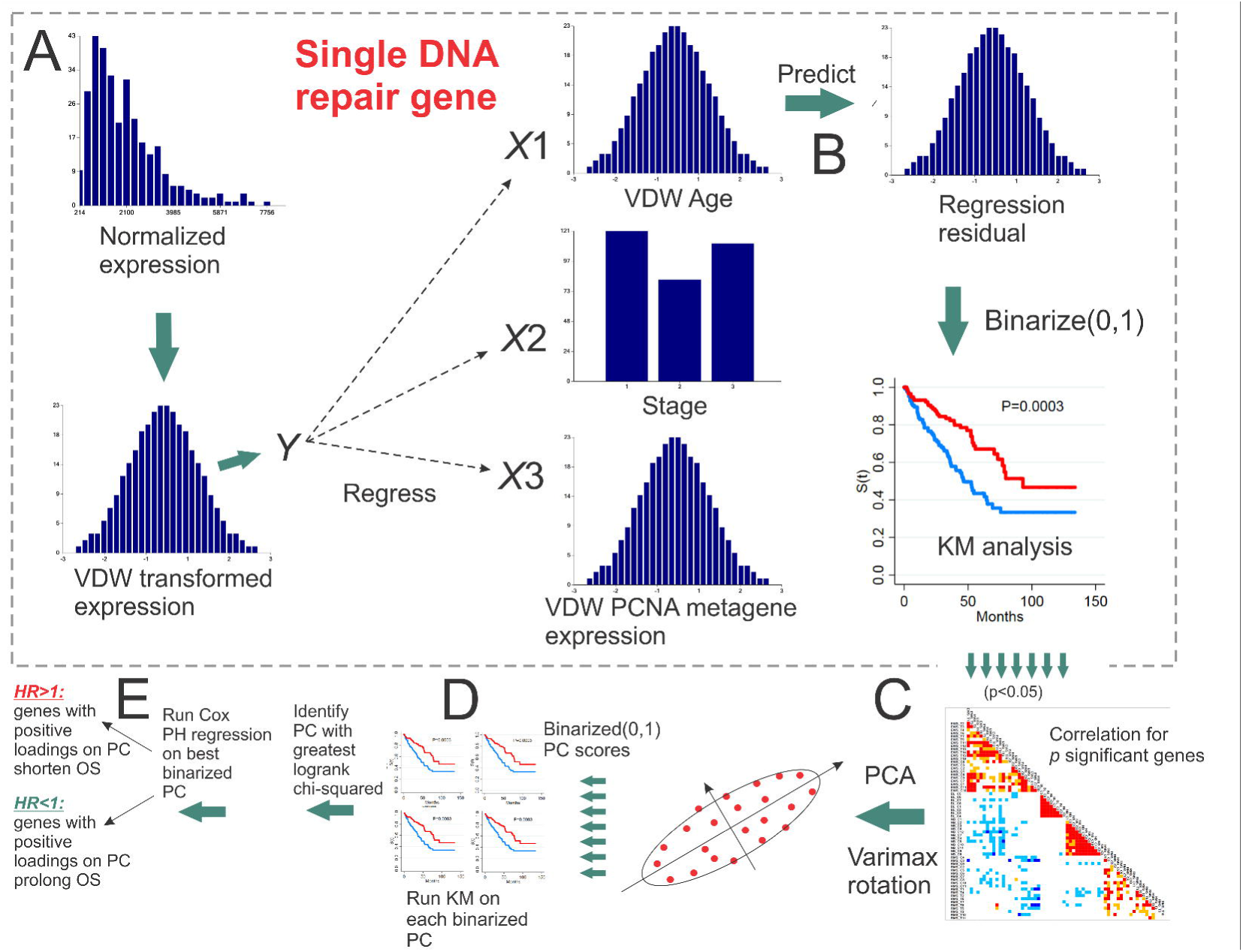
Workflow for identifying the “best binarized PC” for a set of significant genes from univariate Kaplan-Meier analyses. **A.** van der Waerden (VDW) scores of log-transformed expression values for each DNA repair gene are regressed on the VDW scores for age at diagnosis, stage, and the PCNA metagene. **B.** The residual values from each linear fit (i.e., expression with the effect of age at diagnosis, stage, and PCNA metagene removed) are then binarized (0,1) and input as the univariate grouping value during Kaplan-Meier (KM) analysis of overall survival (OS). **C.** PCA with Varimax orthogonal rotation is performed on the correlation matrix of *p* significant binarized residual vectors. **D.** Each of the *p* principal component (PC) score vectors is binarized (0,1) and input into univariate KM analysis. E. The PC resulting in the greatest KM chi-squared value is selected as the “best binarized PC,” and univariate Cox PH regression is then run on the best binarized PC to determine if positive (negative) PC score values are associated with prolonged (shortened) OS.

### Empirical P-value Tests of Survival Using Randomly Selected Genes

For each cancer, the single best binarized PC that resulted in the greatest chi-squared statistic during maximum likelihood KM analysis was considered to the be the “observed” test statistic. Recall, this test statistic for the best binarized PC was initially based on individual DNA repair genes whose adjusted gene expression resulted in a significant KM test. Let the number of significant DNA repair genes for a best binarized PC be *p*. We used *B* = 1, 000 iterations for empirical p-value testing. During each *b*th iteration, a random set of *p* DNA repair genes with the same adjustment to expression were selected, followed by correlation analysis, and then PC extraction via eigendecomposition of the *p* × *p* correlation matrix. Each PC was then binarized and used in KM analysis to determine which PC resulted in the greatest chi-squared test statistic for the set of *p* random genes. After *B* iterations, the empirical p-value was equal to

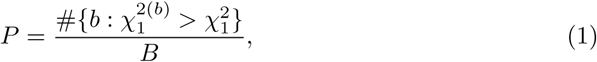

where 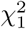 is the “observed” 1 d.f. chi-squared test statistic from ML-based KM analysis based on the best binarized PC, and 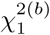 is the chi-squared statistic from the best binarized PC extracted from the correlation matrix of p randomly selected DNA repair genes used in KM analysis during the bth iteration. The bottom of Figure 1 illustrates how the correlation matrix of *p* genes with significant KM analysis were employed to obtain the best binarized PC for predicting OS.

### Genomic Event Rates (GERs)

For each cancer, we also summed the number of pathogenic somatic mutations, deletions, and amplifications in the set of 20 drive genes, and in each of the 8 groups of DNA repair genes (DRR, BER, NHEJ, MMR, TLS, DDS, HRR, NER). The genomic event rate (GER) of each type of event was then determined by dividing the sum by the number of genes in the group and the number of tumors obtained for each of the cancers considered. This led to the GER in units of events per gene-tumor. Hierarchical cluster analysis was then used to cluster values of GER for each cancer. Euclidean distance was used as the distance function, while the unweighted pair group method with arithmetic mean (UPGMA) was used for the agglomeration method.

### Removing Redundant Genes in Gene Lists

*PCNA* itself was a listed as a BER gene, but was removed from the list of DNA repair genes because our primary goal was to remove the genome-wide association of *PCNA* with other genes from the expression of DNA repair genes. For each cancer, any DNA repair gene that was also listed as a cancer-specific driver gene was removed from the list of DNA repair genes, because a driver gene typically has a high frequency of mutations (and other events), and therefore its inclusion in survival analysis and calculation of GERs in sets of DNA repair genes would be counted twice and confound results. DNA repair genes listed multiple times in the repair pathways described above include *POLD1*, *POLE*, *PLOH*, *POLM*, *RECOL2*, *PCNA*, *LIG1*, *BLM*, *BRCA1*, *FANCA*, *FANCC*, and *XPF*. Only the first occurrence of these genes in their respective lists was used. Altogether, a final list of 126 unique (non-redundant) DNA repair genes was constructed and used for all cancers.

## 4 RESULTS

Table 1 lists sample sizes and cancer sites for which Kaplan-Meier logrank tests of OS based on the best binarized PC from correlation of multiple DNA repair gene expression adjusted for age and the *PCNA* metagene. For the simultaneous adjustment of expression by age and *PCNA* metagene (first A,P column), all of the cancers resulted in a significant KM logrank test for the best binarized PC; however, the empirical p-values for random selection of genes resulted in a significant KM test (second A,P column) for 3 cancers, namely, AML, bladder, and sarcoma.

**Table 1.**
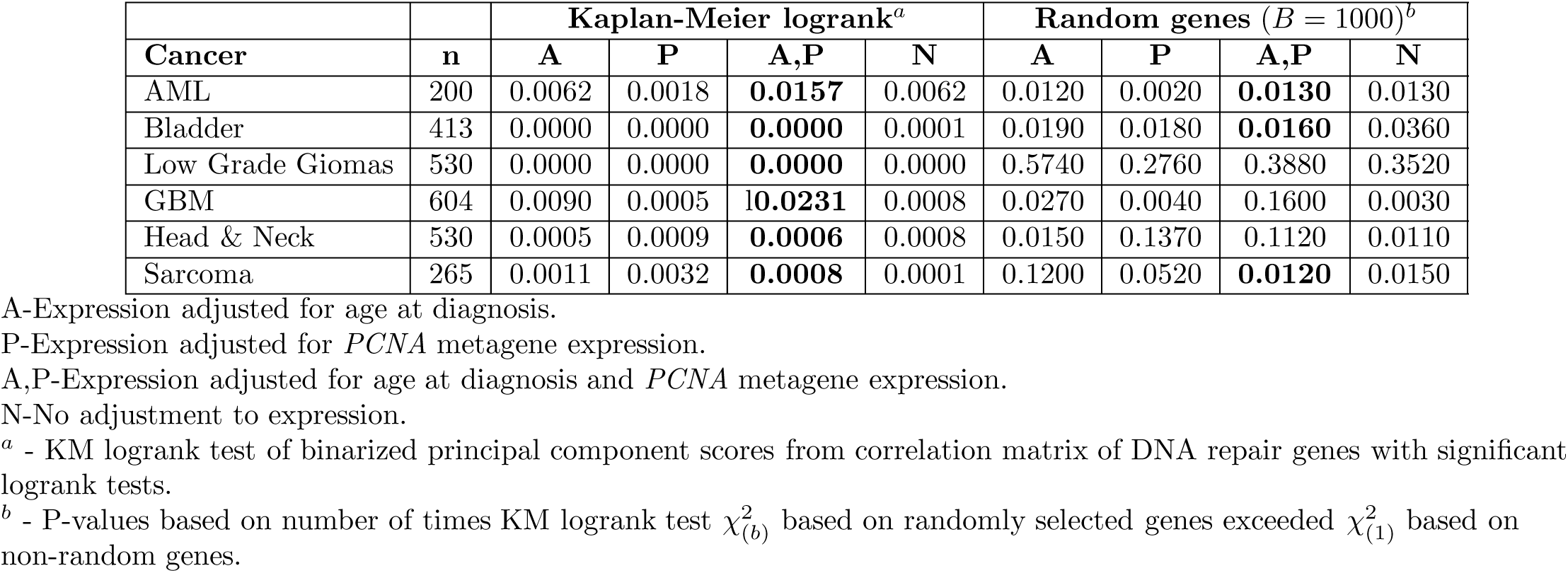
Kaplan-Meier logrank test p-values for best principal component (best binarized PC) derived from correlation matrix of DNA repair genes with significant individual KM tests after adjustment of expression for age and *PCNA* metagene effects. Cancers listed had only age at diagnosis available in TCGA clinical data.

Table 2 lists sample sizes and cancer sites for KM tests performed on the best binarized PC for multiple DNA repair gene expression adjusted for age, stage, and the *PCNA* metagene. For the simultaneous adjustment of expression for age, stage, and *PCNA* metagene (first A,S,P column), 11 of the 12 cancer resulted in PCs whose KM test result were significant. However, when random gene sets of the same size were selected, (second A,S,P column), only 8 of the 12 cancers were significant: breast, colorectal, liver, lung, lung squamous cell, melanoma, renal papillary cancer, and stomach. The combined results in Tables 1 and 2 suggest that while age, stage, and *PCNA* metagene adjusted expression of DNA repair gene expression resulted in significant KM tests for 94% of the cancers (17/18 tests), the selection of random gene sets of the same size resulted in 61% (11/18) cancers showing significant prediction of OS by adjusted DNA repair gene expression.

**Table 2.** Kaplan-Meier logrank test p-values for best principal component (best binarized PC) derived from correlation matrix of DNA repair genes with significant individual KM tests after adjustment of expression for age, stage, and *PCNA* metagene effects. Cancer listed had both age at diagnosis and stage available in TCGA clinical data.

| Cancer | n | Kaplan-Meier logrank<sup>a</sup> | | | | | | Random genes ($B = 1000$)<sup>b</sup> | | | | | |
| --- | --- | --- | --- | --- | --- | --- | --- | --- | --- | --- | --- | --- | --- |
|  |  | A | S | P | A,S | A,S,P | N | A | S | A,S | P | A,S,P | N |
| Breast | 1105 | 0.0057 | 0.0049 | 0.0004 | 0.0069 | **0.0015** | 0.0060 | 0.0790 | 0.0450 | 0.0050 | 0.0860 | **0.0350** | 0.0650 |
| Cervical | 309 | 0.0071 | 0.0014 | 0.0015 | 0.0100 | **0.0151** | 0.0088 | 0.2150 | 0.0630 | 0.0120 | 0.2530 | 0.0730 | 0.2810 |
| Colorectal | 633 | 0.0550 | 0.0255 | 0.0048 | 0.0188 | **0.0106** | 0.0330 | 0.2050 | 0.0700 | 0.0120 | 0.0450 | **0.0340** | 0.1160 |
| Liver | 379 | 0.0000 | 0.0002 | 0.0004 | 0.0001 | **0.0027** | 0.0001 | 0.1150 | 0.2220 | 0.0020 | 0.2140 | **0.0130** | 0.1720 |
| Lung | 522 | 0.0120 | 0.0008 | 0.0003 | 0.0021 | **0.0036** | 0.0070 | 0.4850 | 0.0560 | 0.0070 | 0.1030 | **0.0400** | 0.3570 |
| Lung Sq. Cell | 505 | 0.0057 | 0.0050 | 0.0040 | 0.0050 | **0.0040** | 0.0057 | 0.0400 | 0.0290 | 0.0110 | 0.0290 | **0.0100** | 0.0330 |
| Ovarian | 603 | 0.0259 | 0.0088 | 0.0184 | 0.0722 | **0.0100** | 0.0183 | 0.2310 | 0.1250 | 0.2010 | 0.5240 | 0.0700 | 0.2060 |
| Melanoma | 479 | 0.0000 | 0.0001 | 0.0002 | 0.0000 | **0.0001** | 0.0001 | 0.0170 | 0.0710 | 0.0190 | 0.0250 | **0.0040** | 0.0670 |
| Renal Clear Cell | 538 | 0.0000 | 0.0000 | 0.0000 | 0.0000 | **0.0003** | 0.0000 | 0.2570 | 0.4230 | 0.0100 | 0.1340 | 0.2780 | 0.4710 |
| Renal Papillary | 292 | 0.0000 | 0.0001 | 0.0011 | 0.0000 | **0.0012** | 0.0000 | 0.0770 | 0.0100 | 0.1060 | 0.0040 | **0.0190** | 0.0320 |
| Stomach | 478 | 0.6412 | 0.0103 | 0.0180 | 0.0103 | 0.0107 | 0.6412 | 0.6570 | 0.0070 | 0.0300 | 0.0060 | **0.0160** | 0.6780 |
| Uterine | 548 | 0.0013 | 0.0024 | 0.0150 | 0.0010 | **0.0072** | 0.0009 | 0.0250 | 0.0390 | 0.3620 | 0.0070 | 0.1070 | 0.0190 |

A-Expression adjusted for age at diagnosis.
S-Expression adjusted for stage.
P-Expression adjusted for *PCNA* metagene expression.
A,S-Expression adjusted for age at diagnosis and stage.
A,S,P-Expression adjusted for age at diagnosis, stage, and *PCNA* metagene expression.
N-No adjustment to expression.
<sup>a</sup> - KM logrank test of binarized principal component scores from correlation matrix of DNA repair genes with significant logrank tests.
<sup>b</sup> - P-values based on number of times KM test $\chi^2_{(b)}$ based on random genes exceeded $\chi^2_{(1)}$ based on non-random genes.

Table 3 lists the DNA repair genes of the best binarized PC for the 11 cancers showing empirical p-values less than 0.05 when adjusting for age, stage, and *PCNA* metagene. When considering the composite of all the genes which were significant survival predictors, pathway activation results indicate upregulation of the BRCA1 pathway, NHEJ pathway, BER pathway, MMR pathway, and downregulation of the NER pathway. ATM signaling (*ATM*, *RAD17*, *RAD50*) was downregulated due to strong downregulation of *ATM*, which has been observed to be due to promoter hypermethylation [11]. With downregulation of the tumor suppressor kinase *ATM* and the checkpoint kinase *CHEK2*, it follows that the ATM intraphase checkpoint would also likely be inactivated. Cells deficient in *ATM* have also been observed to not reduce transcription following DSBs, a phenotype which has been called “radiosensitized DNA synthesis.” In addition, ataxia-telangiaectasia (A-T) cells deficient in ATM are known to repair DSBs following exposure to IR [12]. With regard to the BRCA1 pathway, Complex B was highly upregulated with downstream upregulation of and G1/S-Phase as well as the HRR pathway (*BLM, BRCA1, MSH2, MS62*, and *RFC*). In addition, *FANCD2* was upregulated, which activates S-Phase checkpoint control. However, Complex C was downregulated (mostly due to *RAD50*), possibly suggesting downregulation of downstream G2/M Phase. In the NHEJ pathway, *ATM*, the cross-linking enzyme Artemis (*DLCRE1C*), and *RAD50* were downregulated, however, *LIG3*, *LIG4*, *NBN*, *PRKDC*, *XRCC1*, *XRCC2*, and *XRCC5* were upregulated. Activated genes in the BER pathway included *POLB*, *POLE*, *XRCC1*, *LIG1*, *LIG3*, and *FEN1*. The MMR pathway activated genes were *EXO1*, *FEN1*, *MSH2*, *MSH6*, *RFC2*, and *RFC3*. Downregulated genes in the NER pathway were the sensitizer *APEX1* and the DNA glycosylate *OGG1*. The NER pathway was mostly downregulated due to downregulation of *HR23B*, *TFIH*, *XPC*, and *XPG*. We also noted that the single gene repair mechanism via *MGMT* was also downregulated, which would increase SSBs and their conversion to DSBs at replication forks [13]. Another observation was that neither *PARP1* or *PARP2* were listed in any of the lists of significant genes, which may indicate co-regulation with *PCNA*, whose effect on expression of other DNA repair genes was removed.

**Table 3.** DNA Repair genes whose upregulation or downregulation prolongs overall survival (OS) for subjects with RNA-Seq data in TCGA. Cancers listed had significant empirical p-value test results (*P* < 0.05).

| Cancer | Upregulation prolongs OS | Downregulation prolongs OS |
| --- | --- | --- |
| AML* | *POLN* | *RAD23A*, *APEX2*, *EME2* |
| Bladder* | *BLM*, *RAD9A*, *MGMT*, *LIG1*, *MUTYH*, *DDB1*, *ERCC5*, *XPC*, *FANCD2*, *MSH5*, *DLCRE1C*, *REV1* | *FANCC*, *ALKBH2*, *APEX2*, *LIG3*, *POLB*, *GTF2H5*, *PMS1*, *PRKDC*, *REV3L* |
| Sarcoma* | *MNAT1*, *APEX1*, *APTX*, *FEN1*, *NEIL3*, *DDB1*, *GTF2H3*, *FANCI*, *PRKDC* | *DLCRE1B*, *POLL*, *CUL4A*, *ERCC2*, *MSH2*, *FANCG* |
| Breast | *RAD50*, *PMS1* | *ATRIP*, *FANCC*, *RAD1*, *RFC3*, *NEIL3*, *EXO1*, *FANCB*, *FANCD2*, *FANCI*, *RAD51*, *XRCC4* |
| Colorectal | *DLCRE1C* | *RAD23A*, *RFC2*, *POLL*, *MLH3*, *FANCL* |
| Renal Papillary | *RAD17*, *OGG1*, *DDB2*, *ERCC2* | *BLM*, *RAD1*, *FEN1*, *LIG1*, *EXO1*, *MSH6*, *BRCA2*, *EME1*, *FANCB*, *LIG4* |
| Lung | *RAD17*, *ALKBH3*, *MGMT*, *MPG*, *NEIL1*, *XPC*, *LIG4*, *POLK*, *REV3L* | *BRCA1*, *NBN*, *RAD1*, *NEIL3*, *MMS19*, *FANCI*, *XRCC5* |
| Lung Sq. Cell | *CHEK2*, *MNAT1*, *APTX*, *TDG*, *FANCE*, *FANCL* | *XRCC1* |
| Melanoma | *ATM*, *MNAT1*, *MBD4*, *NEIL1*, *ERCC5*, *RAD23B*, *DLCRE1C* | *MDC1*, *NBN*, *MUTYH*, *POLE*, *UNG*, *FANCE*, *FANCI*, *POLI*, *POLK* |
| Stomach | *CUL4A*, *POLQ* |  |

\*Stage not available, expression adjusted only by age and *PCNA* metagene.

Results of cluster analysis of the GER for various cancers is shown in Figure 2. A total of 4 clusters of cancers were discernible in the data. In spite of all the cancers exhibiting high GERs for driver mutations, cancers in cluster 1 portrayed strong upregulation of genomic amplification in DNA repair genes, while cancers in cluster 2 reveal downregulation of amplification, deletion, and mutation in DNA genes. Melanoma and ovarian cancer clustered furthest away from the previously described clusters, mostly because of the unique patterns among GERs which emerged. Regarding driver genes, both melanoma and ovarian cancer exhibited greater rates of amplifications, but had lower rates of deletions. Additionally, while melanoma revealed increased rates of mutations in DNA repair genes and decreased rates of deletions in DNA repairs genes, ovarian cancer showed the opposite pattern, with lower rates of mutations in DNA repair genes and greater rates of deletions in DNA repair genes. Table 4 lists qualitative patterns which emerged from the cluster analysis of GERs shown in Figure 2.

**Fig 2.**
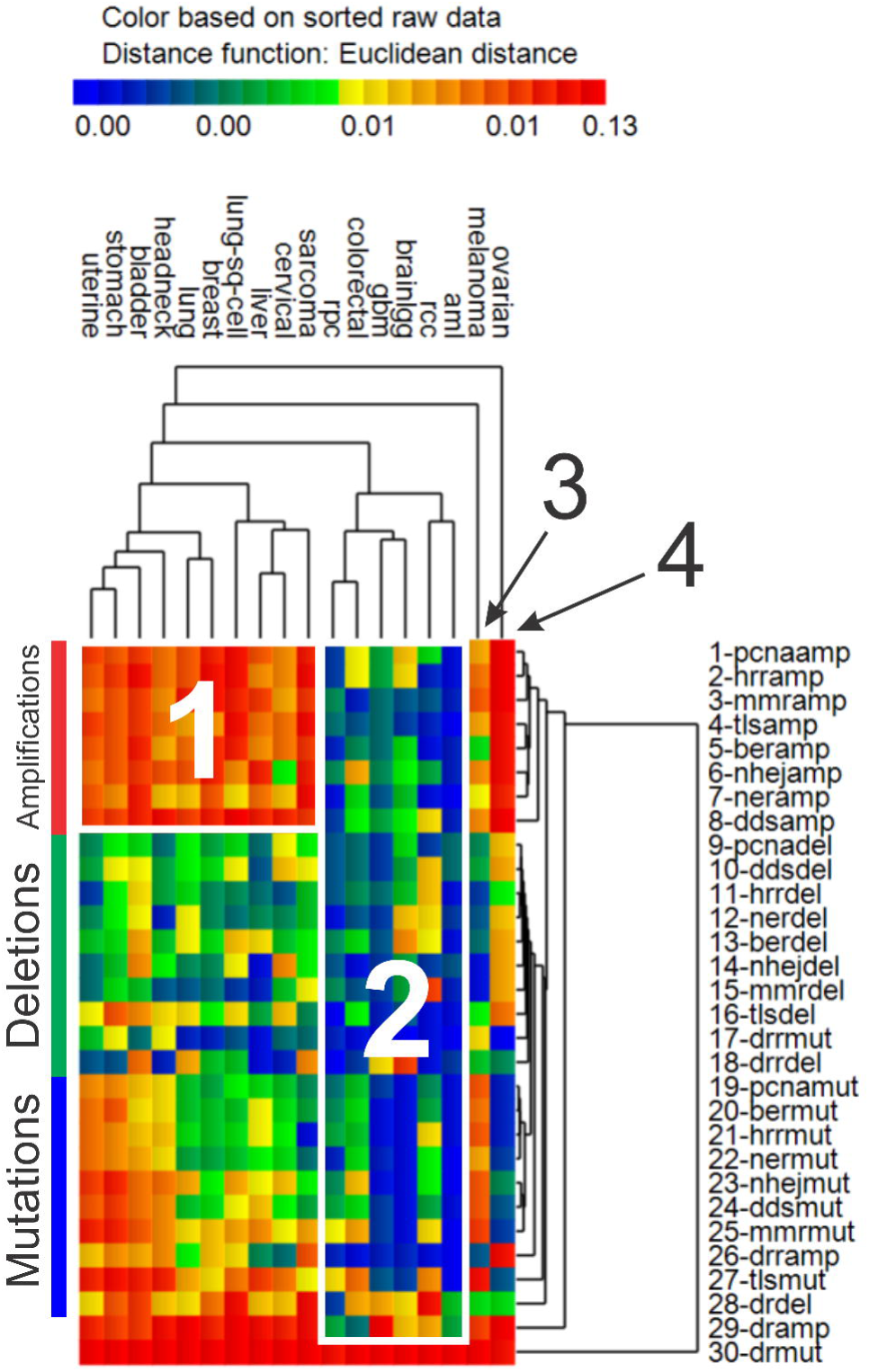
Genomic event rates. Hierarchical cluster analysis results of genomic event rates (GER, per tumor-gene) for somatic mutations (“mut”), deletions (“del”), and amplifications (“amp”) in driver genes (“dr”) and the 8 groups of DNA repair genes (DRR, BER, NHEJ, MMR, TLS, DDS, HRR, NER). Euclidean distance used as the distance function, with unweighted pair group method with arithmetic mean (UPGMA) as the agglomeration method.

**Table 4.**
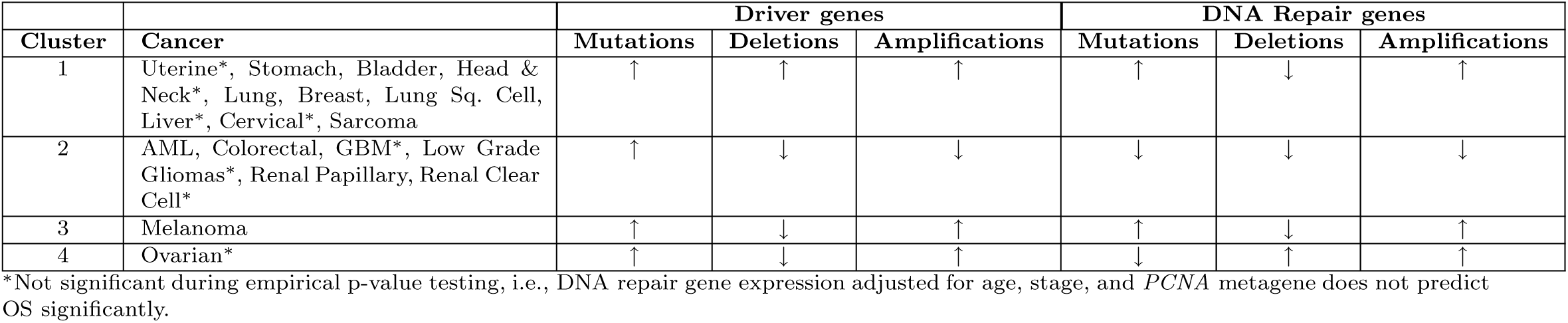
Qualitative patterns of genomic event rates (GER) per gene-tumor for each cluster identified during hierarchical cluster analysis (from Figure 2). Opportunistic cancers for further study in cluster 2 are AML, Colorectal, and Renal Papillary.

Figures S1-S36 illustrate for each cancer investigated a KM plot and density (pdf) plot of p-values during random selection of gene sets for the best binarized PCs derived from sets of genes, for which each gene had its own significant KM test after the various adjustments for age, stage, and *PCNA* metagene. P-values listed in Tables 1 and 2 were extracted from Figures S1-S36.

## 5 DISCUSSION

Cancer is a multifactorial disease which depends on a constellation of factors involving genomic instability, selective genetic pressure from somatic mutations and polymorphisms, and gene-environment interactions. Two important hallmarks of cancer are the persistent high level of somatic mutations in driver genes and genome-wide error-prone transcriptional coding by polymerases. Together, these mechanisms directly and indirectly abrogate apoptosis, leading to a growth advantage and prolonged survival. The growing mutational load that ensues is countered by DNA repair mechanisms with a high level of fidelity. As the costs of nextgen DNA sequencing and RNA-Seq analysis decrease, there will continue to be new information available regarding the balance between mutational load and DNA repair.

Our approach employed two levels of statistical testing, one that merely involved straightforward ML estimation and another based on random gene selection, which resulted in empirical p-values. The ML-based survival prediction with adjusted DNA repair gene expression was significant for most of the cancers; however, survival prediction based on empirical p-values was significant for fewer cancers. It is now known that identification of sets of genes from genome-wide annotation lists will result in false positives that are associated with the *PCNA* metagene. Our focus was to specifically target DNA repair gene expression, remove the effect of the *PCNA* metagene, age at diagnosis, and stage, to determine if significant lists can be obtained. Not surprisingly, after the adjustments, many of the cancers revealed DNA repair genes which significantly predicted OS.

Pathway analysis results indicate a pattern suggestive of downregulation of primary damage signaling kinase (*ATM*) and initial BER pathway components (*APEX1* and *OGG1*), and when combined with increased pathogenic somatic mutations in driver genes (e.g., *TP53*), our results may indicate that initial damage signaling mechanisms are abrogated, but the full compliment of DNA repair is intact. There was also an observation that a single-gene repair mechanism was abrogated, which may be suggestive that minor repair mechanisms are also abrogated.

We also assessed GERs of the various cancers, and confirmed that all of the cancers had high somatic mutation rates in driver genes. There were also two main clusters of cancers identified, which portrayed either high levels of amplification in DNA repair genes or low GERs for mutations, deletions, and amplification in DNA repair genes. The latter group of cancers including AML, colorectal, GBM, low grade gliomas, and renal papillary (cluster 2 in Table 4), may be more opportunistic for investigating further because the low levels of mutations, deletions, and amplification in DNA repair genes would suggest that the suppressed and activated DNA repair pathways in these cancers are intact and unaltered genomically. When considering results of empirical p-value testing, this list of cancers becomes reduced to AML, colorectal, and renal papillary, since OS for GBM and low grade gliomas could not be predicted significantly using random gene tests.

Implementing our recommendations for therapy would require a way to actively suppress or activate various DNA repair pathways, depending on the pattern of repair observed within certain cancers. RNA interference (RNAi) has recently proven to be a valid mechanism for engaging RNAi pathways for the purpose of targeted therapy for cancer patients [14]. Recently, it has been shown that silencing RNA (siRNA) administered systemically to a human can produce a specific gene inhibition (reduction in mRNA and protein) by an RNAi mechanism of action [15]. The advantages of RNAi therapy do not currently outweigh the disadvantages, because of the large uncertainties surrounding the efficacy, delivery efficiency, pharmacokinetics, on-target, and off-target activity on other pathways.

The translational value of our results are established by the potential of novel patterns of DNA repair gene expression in cancer, which could prove useful in animal studies, transgenics, and xenograft models, etc., in order to understand if suppression (inhibition or knock-down) or activation (overexpression) of the genes identified inhibit tumor growth and improve survival [16–18]. Adjustment of DNA repair gene expression by the *PCNA* metagene has enabled us to view cancer from a distant perspective based on high-granularity involvement of DNA repair pathways in cancer. This view will hopefully enforce an appreciation among biologists and oncologists for the translational value of pursuing experimental inhibition and overexpression studies, as well as randomized control trials for establishing safety and evaluating efficacy.

We did not comparatively assess numerous techniques for their computational efficiency, scalability, or differences in OS survival prediction. We also did not evaluate differences between using progression free survival (PFS) vs. OS, or bootstrapping effects on results. An advantage of using OS instead of PFS with TCGA data, is that there are fewer late stage (i.e., stage 4) cancers in TCGA data. In fact, prostate cancer tumors in TCGA have much more data regarding PFS than OS. As such, OS will likely be more informative for more virulent tumors. The work presented here suggests that investigation of the effects of *PNCA* metagene on DNA repair gene expression and OS survival prediction can establish new leads for future research on cancer therapeutics.

There are several challenging issues surrounding OS prediction using TCGA data. First, there is the problem of unknown upstream effects of germline polymorphisms and DNA repair deficiencies which may result in a variety of unknown influences. Second, the difficulty presented by cellular niching and high levels of clonal heterogeneity in tumors presents a challenge for fully unraveling the associations observed in this study. The TCGA data used are not based on single-cell RNA-Seq analysis, which would be helpful for elucidating heterogeneity effects; however, the large variation in genotypes would exacerbate the present uncertainties surrounding our attempt to portray the role of DNA repair genes in cancer survival. We also did not consider DNA microsatellite instability, methylation status, or chromosome aberrations, which would overlay more complexity on the models developed.

In conclusion, our hypothesis-driven focus to target DNA repair gene expression adjusted for the *PCNA* metagene as a means of predicting OS in various cancers resulted in statistically significant sets of DNA repair genes. We also identified that AML, colorectal, and renal papillary cancers may be more opportunistic for future knowledge discovery because of the low rates of mutations, deletions, and amplification in DNA repair genes which predict OS in these cancers. The most opportunistic cancer appears to be AML, since it harbors the lowest rates of somatic mutations, deletions, and amplifications in DNA repair genes.

## Supporting information

S1 Fig. Acute Myelogenous Leukemia: Kaplan-Meier logrank test results for single best binarized principal component (0,1) extracted from correlation matrix for *p* genes whose adjusted expression (via age, *PCNA* metagene) resulted in significant gene-specific KM logrank tests.

S2 Fig. Acute Myelogenous Leukemia: Empirical p-value test results for single best binarized principal component (0,1) extracted from correlation matrix for *p* genes whose adjusted expression (via age, *PCNA* metagene) resulted in significant gene-specific KM logrank tests. Square symbols denote the observed −log(*P*) for the best binarized PC based on maximum likelihood analysis using KM analysis. Kernel density curves reflect the distribution of −log(*P*) for KM analysis of the best binarized PC using p randomly selected genes *B* = 1000 times.

S3 Fig. Bladder: Kaplan-Meier logrank test results for single best binarized principal component (0,1) extracted from correlation matrix for *p* genes whose adjusted expression (via age, *PCNA* metagene) resulted in significant gene-specific KM logrank tests.

S4 Fig. Bladder: Empirical p-value test results for single best binarized principal component (0,1) extracted from correlation matrix for *p* genes whose adjusted expression (via age, *PCNA* metagene) resulted in significant gene-specific KM logrank tests. Square symbols denote the observed −log(*P*) for the best binarized PC based on maximum likelihood analysis using KM analysis. Kernel density curves reflect the distribution of −log(*P*) for KM analysis of the best binarized PC using *p* randomly selected genes B = 1000 times.

S5 Fig. Breast: Kaplan-Meier logrank test results for single best binarized principal component (0,1) extracted from correlation matrix for *p* genes whose adjusted expression (via age, stage, *PCNA* metagene) resulted in significant gene-specific KM logrank tests.

S6 Fig. Breast: Empirical p-value test results for single best binarized principal component (0,1) extracted from correlation matrix for *p* genes whose adjusted expression (via age, stage, *PCNA* metagene) resulted in significant gene-specific KM logrank tests. Square symbols denote the observed −log(*P*) for the best binarized PC based on maximum likelihood analysis using KM analysis. Kernel density curves reflect the distribution of −log(*P*) for KM analysis of the best binarized PC using *p* randomly selected genes B = 1000 times.

S7 Fig. Cervical: Kaplan-Meier logrank test results for single best binarized principal component (0,1) extracted from correlation matrix for *p* genes whose adjusted expression (via age, stage, *PCNA* metagene) resulted in significant gene-specific KM logrank tests.

S8 Fig. Cervical: Empirical p-value test results for single best binarized principal component (0,1) extracted from correlation matrix for *p* genes whose adjusted expression (via age, stage, *PCNA* metagene) resulted in significant gene-specific KM logrank tests. Square symbols denote the observed −log(*P*) for the best binarized PC based on maximum likelihood analysis using KM analysis. Kernel density curves reflect the distribution of −log(*P*) for KM analysis of the best binarized PC using *p* randomly selected genes B = 1000 times.

S9 Fig. Colorectal: Kaplan-Meier logrank test results for single best binarized principal component (0,1) extracted from correlation matrix for *p* genes whose adjusted expression (via age, stage, *PCNA* metagene) resulted in significant gene-specific KM logrank tests.

S10 Fig. Colorectal: Empirical p-value test results for single best binarized principal component (0,1) extracted from correlation matrix for *p* genes whose adjusted expression (via age, stage, *PCNA* metagene) resulted in significant gene-specific KM logrank tests. Square symbols denote the observed −log(*P*) for the best binarized PC based on maximum likelihood analysis using KM analysis. Kernel density curves reflect the distribution of −log(*P*) for KM analysis of the best binarized PC using *p* randomly selected genes B = 1000 times.

S11 Fig. Glioblastoma Multiforme: Kaplan-Meier logrank test results for single best binarized principal component (0,1) extracted from correlation matrix for *p* genes whose adjusted expression (via age, *PCNA* metagene) resulted in significant gene-specific KM logrank tests.

S12 Fig. Glioblastoma Multiforme: Empirical p-value test results for single best binarized principal component (0,1) extracted from correlation matrix for *p* genes whose adjusted expression (via age, *PCNA* metagene) resulted in significant gene-specific KM logrank tests. Square symbols denote the observed −log(*P*) for the best binarized PC based on maximum likelihood analysis using KM analysis. Kernel density curves reflect the distribution of −log(*P*) for KM analysis of the best binarized PC using *p* randomly selected genes *B* = 1000 times.

S13 Fig. Head and Neck: Kaplan-Meier logrank test results for single best binarized principal component (0,1) extracted from correlation matrix for *p* genes whose adjusted expression (via age, *PCNA* metagene) resulted in significant gene-specific KM logrank tests.

S14 Fig. Head and Neck: Empirical p-value test results for single best binarized principal component (0,1) extracted from correlation matrix for *p* genes whose adjusted expression (via age, *PCNA* metagene) resulted in significant gene-specific KM logrank tests. Square symbols denote the observed −log(*P*) for the best binarized PC based on maximum likelihood analysis using KM analysis. Kernel density curves reflect the distribution of −log(*P*) for KM analysis of the best binarized PC using *p* randomly selected genes *B* = 1000 times.

S15 Fig. Low Grade Gliomas: Kaplan-Meier logrank test results for single best binarized principal component (0,1) extracted from correlation matrix for *p* genes whose adjusted expression (via age, *PCNA* metagene) resulted in significant gene-specific KM logrank tests.

S16 Fig. Low Grade Gliomas: Empirical p-value test results for single best binarized principal component (0,1) extracted from correlation matrix for *p* genes whose adjusted expression (via age, *PCNA* metagene) resulted in significant gene-specific KM logrank tests. Square symbols denote the observed −log(*P*) for the best binarized PC based on maximum likelihood analysis using KM analysis. Kernel density curves reflect the distribution of −log(*P*) for KM analysis of the best binarized PC using *p* randomly selected genes *B* = 1000 times.

S17 Fig. Liver: Kaplan-Meier logrank test results for single best binarized principal component (0,1) extracted from correlation matrix for *p* genes whose adjusted expression (via age, stage, *PCNA* metagene) resulted in significant gene-specific KM logrank tests.

S18 Fig. Liver: Empirical p-value test results for single best binarized principal component (0,1) extracted from correlation matrix for *p* genes whose adjusted expression (via age, stage, *PCNA* metagene) resulted in significant gene-specific KM logrank tests. Square symbols denote the observed −log(*P*) for the best binarized PC based on maximum likelihood analysis using KM analysis. Kernel density curves reflect the distribution of −log(*P*) for KM analysis of the best binarized PC using *p* randomly selected genes *B* = 1000 times.

S19 Fig. Lung Sq. Cell: Kaplan-Meier logrank test results for single best binarized principal component (0,1) extracted from correlation matrix for *p* genes whose adjusted expression (via age, stage, *PCNA* metagene) resulted in significant gene-specific KM logrank tests.

S20 Fig. Lung Sq. Cell: Empirical p-value test results for single best binarized principal component (0,1) extracted from correlation matrix for *p* genes whose adjusted expression (via age, stage, *PCNA* metagene) resulted in significant gene-specific KM logrank tests. Square symbols denote the observed −log(*P*) for the best binarized PC based on maximum likelihood analysis using KM analysis. Kernel density curves reflect the distribution of −log(*P*) for KM analysis of the best binarized PC using *p* randomly selected genes *B* = 1000 times.

S21 Fig. Lung: Kaplan-Meier logrank test results for single best binarized principal component (0,1) extracted from correlation matrix for *p* genes whose adjusted expression (via age, stage, *PCNA* metagene) resulted in significant gene-specific KM logrank tests.

S22 Fig. Lung: Empirical p-value test results for single best binarized principal component (0,1) extracted from correlation matrix for *p* genes whose adjusted expression (via age, stage, *PCNA* metagene) resulted in significant gene-specific KM logrank tests. Square symbols denote the observed −log(*P*) for the best binarized PC based on maximum likelihood analysis using KM analysis. Kernel density curves reflect the distribution of −log(*P*) for KM analysis of the best binarized PC using *p* randomly selected genes *B* = 1000 times.

S23 Fig. Melanoma: Kaplan-Meier logrank test results for single best binarized principal component (0,1) extracted from correlation matrix for *p* genes whose adjusted expression (via age, stage, *PCNA* metagene) resulted in significant gene-specific KM logrank tests.

S24 Fig. Melanoma: Empirical p-value test results for single best binarized principal component (0,1) extracted from correlation matrix for *p* genes whose adjusted expression (via age, stage, *PCNA* metagene) resulted in significant gene-specific KM logrank tests. Square symbols denote the observed −log(*P*) for the best binarized PC based on maximum likelihood analysis using KM analysis. Kernel density curves reflect the distribution of −log(*P*) for KM analysis of the best binarized PC using *p* randomly selected genes *B* = 1000 times.

S25 Fig. Ovarian: Kaplan-Meier logrank test results for single best binarized principal component (0,1) extracted from correlation matrix for *p* genes whose adjusted expression (via age, stage, *PCNA* metagene) resulted in significant gene-specific KM logrank tests.

S26 Fig. Ovarian: Empirical p-value test results for single best binarized principal component (0,1) extracted from correlation matrix for *p* genes whose adjusted expression (via age, stage, *PCNA* metagene) resulted in significant gene-specific KM logrank tests. Square symbols denote the observed −log(*P*) for the best binarized PC based on maximum likelihood analysis using KM analysis. Kernel density curves reflect the distribution of −log(*P*) for KM analysis of the best binarized PC using *p* randomly selected genes *B* = 1000 times.

S27 Fig. Renal Clear Cell: Kaplan-Meier logrank test results for single best binarized principal component (0,1) extracted from correlation matrix for *p* genes whose adjusted expression (via age, stage, *PCNA* metagene) resulted in significant gene-specific KM logrank tests.

S28 Fig. Renal Clear Cell: Empirical p-value test results for single best binarized principal component (0,1) extracted from correlation matrix for *p* genes whose adjusted expression (via age, stage, *PCNA* metagene) resulted in significant gene-specific KM logrank tests. Square symbols denote the observed −log(*P*) for the best binarized PC based on maximum likelihood analysis using KM analysis. Kernel density curves reflect the distribution of −log(*P*) for KM analysis of the best binarized PC using *p* randomly selected genes *B* = 1000 times.

S29 Fig. Renal Papillary: Kaplan-Meier logrank test results for single best binarized principal component (0,1) extracted from correlation matrix for *p* genes whose adjusted expression (via age, stage, *PCNA* metagene) resulted in significant gene-specific KM logrank tests.

S30 Fig. Renal Papillary: Empirical p-value test results for single best binarized principal component (0,1) extracted from correlation matrix for *p* genes whose adjusted expression (via age, stage, *PCNA* metagene) resulted in significant gene-specific KM logrank tests. Square symbols denote the observed −log(*P*) for the best binarized PC based on maximum likelihood analysis using KM analysis. Kernel density curves reflect the distribution of −log(*P*) for KM analysis of the best binarized PC using *p* randomly selected genes *B* = 1000 times.

S31 Fig. Sarcoma: Kaplan-Meier logrank test results for single best binarized principal component (0,1) extracted from correlation matrix for *p* genes whose adjusted expression (via age, stage, *PCNA* metagene) resulted in significant gene-specific KM logrank tests.

S32 Fig. Sarcoma: Empirical p-value test results for single best binarized principal component (0,1) extracted from correlation matrix for *p* genes whose adjusted expression (via age, *PCNA* metagene) resulted in significant gene-specific KM logrank tests. Square symbols denote the observed −log(*P*) for the best binarized PC based on maximum likelihood analysis using KM analysis. Kernel density curves reflect the distribution of −log(*P*) for KM analysis of the best binarized PC using *p* randomly selected genes *B* = 1000 times.

S33 Fig. Stomach: Kaplan-Meier logrank test results for single best binarized principal component (0,1) extracted from correlation matrix for *p* genes whose adjusted expression (via age, stage, *PCNA* metagene) resulted in significant gene-specific KM logrank tests.

S34 Fig. Stomach: Empirical p-value test results for single best binarized principal component (0,1) extracted from correlation matrix for *p* genes whose adjusted expression (via age, stage, *PCNA* metagene) resulted in significant gene-specific KM logrank tests. Square symbols denote the observed −log(*P*) for the best binarized PC based on maximum likelihood analysis using KM analysis. Kernel density curves reflect the distribution of −log(*P*) for KM analysis of the best binarized PC using *p* randomly selected genes *B* = 1000 times.

S35 Fig. Uterine: Kaplan-Meier logrank test results for single best binarized principal component (0,1) extracted from correlation matrix for *p* genes whose adjusted expression (via age, stage, *PCNA* metagene) resulted in significant gene-specific KM logrank tests.

S36 Fig. Uterine: Empirical p-value test results for single best binarized principal component (0,1) extracted from correlation matrix for *p* genes whose adjusted expression (via age, stage, *PCNA* metagene) resulted in significant gene-specific KM logrank tests. Square symbols denote the observed −log(*P*) for the best binarized PC based on maximum likelihood analysis using KM analysis. Kernel density curves reflect the distribution of −log(*P*) for KM analysis of the best binarized PC using *p* randomly selected genes *B* = 1000 times.

## Acknowledgments

Research was supported by NASA Grant NNX-12AO52A. LEP wrote the manuscript, developed the workflow, and performed the majority of computational modeling, while TK reviewed and confirmed the statistical procedures.

